# First characterization of *Mycobacterium avium* subsp. *paratuberculosis* strain types in cattle from Kushiro Subprefecture, eastern Hokkaido, Japan

**DOI:** 10.64898/2026.04.27.721228

**Authors:** Shigeru Matsuzawa, Masako Narita

## Abstract

Johne’s disease (JD), caused by *Mycobacterium avium* subsp. *paratuberculosis* (MAP), is an important livestock disease in Japan. We typed 48 field isolates from cattle feces collected on 13 farms in Kushiro Subprefecture during 2024–2025 and 62 MAP-positive cattle fecal samples collected in the same region during 2025–2026 using a real-time PCR assay targeting the Type S-specific arylsulfatase gene. No Type S strains were detected among the cultured isolates or fecal samples examined in this study, suggesting that the cattle cases analyzed were more consistent with Type C than with Type S strains. Broader surveys are needed to define MAP strain diversity in Japan and improve control strategies.

---

Paratuberculosis, also known as Johne’s disease (JD), is an important livestock disease in Japan, particularly in dairy and beef cattle. It is caused by *Mycobacterium avium* subsp. *paratuberculosis* (MAP) and is characterized by chronic enteritis. In Japan, rigorous control measures have been implemented, including mandatory periodic testing of cattle used for breeding, at least once every 5 years. As a result, approximately 1,000 positive cases are identified annually, although most affected animals show only mild or no clinical signs [1]. Although the use of advanced diagnostic methods such as ELISA and real-time PCR has improved diagnostic accuracy [1], fecal culture is still considered the gold standard for diagnosing MAP infection [2,6].

The existence of two major MAP lineages, Type C and Type S, is of considerable epidemiological and bacteriological importance. For example, Type S strains grow more poorly than Type C strains on commonly used media such as Herrold’s egg yolk medium (HEYM) [3–6]. Despite this marked difference in culturability, the distribution of Type C and Type S MAP strains in cattle in Japan has not been systematically investigated, particularly using direct testing of MAP-positive fecal materials. Restriction fragment length polymorphism analysis has been widely used to distinguish between the two major MAP groups based on the copy number and genomic locations of insertion sequence (IS) *900* [3] or on highly conserved single-nucleotide polymorphisms within IS*1311* [4]. However, this method has limited discriminatory power [6]. Although whole-genome sequencing provides the highest resolution for strain discrimination, it remains costly and labor-intensive for routine use. More recently, a sensitive assay capable of distinguishing between the two MAP types directly from fecal samples has been developed by targeting a genomic region unique to Type S strains, namely the arylsulfatase gene [6]. In this study, we investigated MAP strain types in cattle from Kushiro Subprefecture by applying a Type S-specific real-time PCR assay to both cultured isolates and MAP-positive fecal samples.

Kushiro Subprefecture in eastern Hokkaido, northern Japan, is characterized by extensive grasslands and substantial cattle and sheep farming. In this study, 48 MAP field isolates recovered from fecal samples collected from cattle diagnosed with Johne’s disease on 13 farms in Kushiro Subprefecture between 2024 and 2025 were used (Table 1). Each isolate represented a different animal, and only a single colony was picked from each culture for downstream analysis. These isolates were cultured on HEYM supplemented with mycobactin (Kyoritsu Seiyaku Co., Tokyo, Japan). After incubation, individual colonies from the slants were suspended in 50 μL of sterile water and boiled for 10 min. The samples were briefly centrifuged, and the supernatant was used as the template. The presence of MAP IS900 was first confirmed using Johne-findpro (FASMAC, Atsugi, Japan) and a LightCycler 480 Real-Time PCR System (Roche, Basel, Switzerland), according to the manufacturer’s instructions. All samples were positive and were subsequently subjected to Type S-specific real-time PCR as previously described [6]. The primer and probe sequences were taken from a previous report [6]: forward, 5′-CACGCTGTTGCGGTATCT-3′; reverse, 5′-CGGTATTTCGCGATCGACT-3′; probe, 5′-FAM-CCGTTCTTC/ZEN/GCCTATCTGCCGTTT-IBFQ-3′. The primers and probe were synthesized by Integrated DNA Technologies (Coralville, IA, USA). A 297-bp synthetic DNA fragment (gBlock™, Integrated DNA Technologies, Coralville, IA, USA) encompassing the target region of the Type S Telford reference genome (accession no. NZ_CP033688.1) was used as a positive control during the initial implementation of the assay. This control was run on a single occasion to verify amplification of the target sequence under our laboratory conditions. Following the initial verification, all subsequent runs were performed under identical reaction conditions using the same instrument. Real-time PCR was performed with minor modifications to the reaction conditions described previously [6], in a final volume of 20 μL containing 10 μL of 2× Probe qPCR mix with UNG (TaKaRa, Kusatsu, Japan), 0.9 μM of each primer, 0.45 μM of probe, 2 μL of template DNA, and nuclease-free water. Reactions were run on a LightCycler 480 Real-Time PCR System. The cycling conditions consisted of a UNG step at 37°C for 10 min, an initial denaturation step at 95°C for 30 s, followed by 35 cycles of 95°C for 15 s and 65°C for 30 s.

**Table 1.** Number of samples used in the current study.

| Farm | No. of field isolates |  | No. of fecal samples |  |
| --- | --- | --- | --- | --- |
|  | dairy cattle | beef cattle | dairy cattle | beef cattle |
| A |  |  |  | 1 |
| B |  |  |  | 1 |
| C |  | 2 |  |  |
| D |  | 3 |  |  |
| E |  | 2 |  |  |
| F | 3 |  | 2 |  |
| G | 3 |  | 1 |  |
| H | 8 |  | 7 |  |
| I | 3 |  |  |  |
| J | 6 |  | 3 |  |
| K | 3 |  |  |  |
| L | 7 |  | 2 |  |
| M | 3 |  |  |  |
| N | 3 |  | 2 |  |
| O | 2 |  | 1 |  |
| P |  |  | 3 |  |
| Q |  |  | 3 |  |
| R |  |  | 2 |  |
| S |  |  | 1 |  |
| T |  |  | 4 |  |
| U |  |  | 3 |  |
| V |  |  | 1 |  |
| W |  |  | 1 |  |
| X |  |  | 6 |  |
| Y |  |  | 2 |  |
| Z |  |  | 1 |  |
| AA |  |  | 1 |  |
| AB |  |  | 4 |  |
| AC |  |  | 1 |  |
| AD |  |  | 1 |  |
| AE |  |  | 7 |  |
| AF |  |  | 1 |  |
|  | 41 | 7 | 60 | 2 |

To address the inherent bias associated with isolate-based sampling on HEYM, an additional 62 fecal samples were included (Table 1). Each fecal sample was obtained from a different animal, and none of these animals overlapped with those from which the 48 cultured isolates were derived. These samples were collected in Kushiro Subprefecture in 2025 and 2026 from cattle that were positive by ELISA and were subsequently diagnosed as Johne’s disease cases according to the Japanese administrative criteria after detection of MAP IS*900* in feces by real-time PCR using Johne-findpro. The animals were then culled in accordance with the control program. Total DNA was extracted from feces using Johne-pure-spin (FASMAC, Atsugi, Japan) in accordance with the manufacturer’s instructions, and the extracted DNA was then subjected to Type S-specific real-time PCR.

In the Type S-specific real-time PCR assay, no positive signals were detected in any of the 48 MAP field isolates or 62 fecal samples. In contrast, all samples tested positive with Johne-findpro, and no apparent PCR inhibition was suggested by the assay’s internal control. All 62 fecal samples had previously been judged MAP-positive by routine diagnostic real-time PCR (Johne-findpro). Although the administrative diagnostic threshold (≥0.001 pg per reaction) is not directly comparable to the analytical limit of detection reported for the Type S-specific assay (0.02 fg per reaction) [6], the consistently negative results in the latter assay provided no evidence for the presence of Type S strains in these materials.

These findings suggest that Type S strains are unlikely to be common among the cattle cases examined in Kushiro Subprefecture of eastern Hokkaido. To the best of our knowledge, no previous study has systematically investigated the distribution of Type C and Type S MAP strains in cattle in Japan. Our findings therefore suggest that the cattle cases examined in Kushiro Subprefecture were more consistent with Type C than with Type S strains. The findings also support the continued use of HEYM-based slants for isolating MAP from cattle in routine diagnostic settings in the study region. This study was limited to cattle from a single subprefecture and did not include sheep or goats, which may differ in the distribution of MAP strain types. In addition, because a Type S-positive control was used only during the initial implementation of the assay, run-to-run analytical performance was not independently verified in every run. Although Type S strains show a strong host preference for sheep, cross-species transmission to cattle can occur and may be more frequent under conditions of close contact [7]. Because the relative proportions of livestock species differ among regions in Japan, broader surveys including other host species are needed to define MAP strain diversity more accurately and to support improved control strategies.

## ACKNOWLEDGMENTS

The authors acknowledge the use of ChatGPT (OpenAI, USA) for assistance with English-language editing and improvement of grammar and readability. The authors reviewed and edited the output and take full responsibility for the content of this manuscript.

## References

1. E. Momotani, “Epidemiological situation and control strategies for paratuberculosis in Japan,” Japanese Journal of Veterinary Research 60 (Supplement) (2012): S19–S29.

2. WOAH, “Chapter 3.1.17. Paratuberculosis (Johne’s disease),” in Manual of Diagnostic Tests and Vaccines for Terrestrial Animals (Paris, France: WOAH, 2021).

3. D. M. Collins, D. M. Gabric, and G. W. de Lisle, “Identification of Two Groups of Mycobacterium paratuberculosis Strains by Restriction Endonuclease Analysis and DNA Hybridization,” Journal of Clinical Microbiology 28, no. 7 (1990): 1591–1596.

4. R. Whittington, I. Marsh, E. Choy and D. Cousins, “Polymorphisms in IS1311, an insertion sequence common to Mycobacterium avium and M. avium subsp. paratuberculosis, can be used to distinguish between and within these species,” Molecular and Cellular Probes 12 (1998): 349–358.

5. R. Mizzi, K. M. Plain, V. J. Timms, I. Marsh and R. J. Whittington, “Characterisation of IS1311 in Mycobacterium avium subspecies paratuberculosis genomes: Typing, continental clustering, microbial evolution and host adaptation,” PLoS ONE 19, no. 2 (2024): e0294570.

6. R. Hodgeman, Y. Liu, S. Rochfort and B. Rodoni, “Development and evaluation of genomics informed real-time PCR assays for the detection and strain typing of Mycobacterium avium subsp. paratuberculosis,” Journal of Applied Microbiology 135 (2024): lxae107.

7. C. Verdugo, E. Pleydell, M. Price-Carter, D. Prattley, D. Collins, G. de Lisle, et al., “Molecular epidemiology of Mycobacterium avium subsp. paratuberculosis isolated from sheep, cattle and deer on New Zealand pastoral farms,” Preventive Veterinary Medicine 117 (2014): 436–446.

